# Phosphorylation of the M1 muscarinic acetylcholine receptor mediates protection in neurodegenerative disease

**DOI:** 10.1101/2021.04.29.440753

**Authors:** Miriam Scarpa, Colin Molloy, Laura Jenkins, Gonzalo Tejeda, Mario Rossi, Louis Dwomoh, Sara Marsango, Zeshan Ahmed, Graeme Milligan, Brian D. Hudson, Andrew B. Tobin, Sophie J. Bradley

## Abstract

There are currently no treatments that can slow the progression of neurodegenerative diseases such as Alzheimer’s disease (AD). There is, however, a growing body of evidence that activation of the M1 muscarinic acetylcholine receptor (M1-receptor) can not only restore memory loss in AD patients, but in preclinical animal models can also slow neurodegenerative disease progression. The generation of an effective medicine targeting the M1-receptor has however been severely hampered by associated cholinergic adverse responses. By using genetically engineered mouse models that express a G protein-biased M1-receptor, we recently established that M1-receptor mediated adverse responses can be minimised by ensuring activating ligands maintain receptor phosphorylation/arrestin-dependent signalling. Here, we use these same genetic models in concert with murine prion disease, a terminal neurodegenerative disease showing key hallmarks of AD, to establish that phosphorylation/arrestin-dependent signalling delivers neuroprotection that both extends normal animal behaviour and prolongs the life span of prion diseased mice. Our data point to an important neuroprotective property inherent to the M1-receptor and indicate that next generation M1-receptor ligands designed to drive receptor phosphorylation/arrestin-dependent signalling would potentially show low adverse responses whilst delivering neuroprotection that will slow disease progression.

## Introduction

The number of people living with dementia, of which Alzheimer’s disease (AD) is the most common form, is estimated to be ∼50 million worldwide (1). This number is predicted to increase to around 130 million by 2050, in line with an aging population. Despite significant efforts to develop disease modifying treatments for AD, there are currently no therapies that can slow or halt disease progression. Symptomatic treatment of memory loss in AD is currently available and delivered by cholinesterase inhibitors that aim to restore defective cholinergic transmission by elevating acetylcholine levels in the brain. These drugs have limited clinical efficacy due mainly to dose-limiting side-effects resulting from the non-selective whole-body up-regulation of cholinergic systems (2, 3). An alternative strategy is to directly activate acetylcholine receptors of the muscarinic family, of which there are five subtypes (M1-M5-receptors). Particular focus has been directed toward the M1-receptor due to high levels of receptor expression in brain regions such as the hippocampus and cortex (4-6), and pro-cognitive effects in pre-clinical animal models (7). Further, clinical trials using the orthosteric agonists, xanomeline and GSK-5, that primarily activate the M1-receptor have shown promising efficacy (8-10). Similarly, M1-receptor selective positive allosteric modulators (PAMs) have been shown to improve cognition in pre-clinical animals models but together with the orthosteric ligands have ultimately failed in the clinic due largely to cholinergic adverse responses, some of which have been ascribed to on-target activity at the M1-receptor (11-13) as well as off-target M2- and M3-receptor activation (14, 15).

To overcome these barriers, we and others have set out to define the optimal pharmacological properties of orthosteric- and allosteric M1-receptor ligands that will deliver clinical efficacy whilst minimising cholinergic adverse responses (16). To this end we have focused our attention on the possible advantages of biased ligands – an approach based on the observation that G protein-coupled receptors (GPCRs) operate by coupling to two fundamental signalling pathways; G protein-dependent signalling and receptor phosphorylation/arrestin-dependent pathways. The promise of ligand-bias is that ligands could be designed to drive GPCR signalling pathways that lead to clinically beneficial outcomes, in preference to ones that result in adverse responses. We have investigated this possibility for the M1-receptor by the generation of a genetically engineered mouse strain that expresses a variant of the M1-receptor where all the intracellular phosphorylation sites have been removed (17). This variant (called M1-PD) is uncoupled from receptor phosphorylation/arrestin-dependent signalling but shows near normal coupling to Gq/11-dependent pathways. By using this receptor variant, we have established that cholinergic-adverse responses to M1-receptor ligands are minimised if receptor phosphorylation/arrestin-dependent signalling is maintained (17). Furthermore, the M1-PD mice have established the importance of M1-receptor phosphorylation/arrestin-dependent signalling in the regulation of anxiety-like behaviours and learning and memory suggesting that maintenance of receptor phosphorylation is important to deliver clinical efficacy as well as minimising adverse responses.

Whereas early studies have provided a framework for the design of M1-receptor ligands for the symptomatic treatment for AD, it has been the emergence of evidence that the M1-receptor might also modify neurodegenerative disease progression that has generated significant attention (7, 18-20). Activation of muscarinic receptors with an orthosteric ligand can regulate the proteolytic processing of amyloid precursor protein thereby reducing the appearance of amyloid β-plaques in a preclinical AD mouse model (19). Our own studies have established that M1-receptor selective PAMs can slow the progression of mouse prion disease thereby maintaining normal animal behaviour and extending life span (18).

Here we have extended these studies by asking if the M1-receptor inherently has neuroprotective properties. By employing the M1-PD mouse strain (17) in combination with mouse prion disease, a progressive terminal neurodegenerative disease that shows many of the hallmarks of human AD (21), we report here that prion disease progresses more rapidly and behavioural abnormalities appear at earlier times in M1-PD mice compared to wild-type controls. The rapid disease onset in M1-PD is further evident in the elevation of neuroinflammatory pathways, including activation of astrocytes and microglia, and the up-regulation of markers of neurodegenerative disease. We conclude that the neuroprotective property of the M1-receptor might be harnessed by next generation M1-receptor drugs for the treatment of AD. M1-receptor-selective drugs could be designed to promote receptor signalling via phosphorylation/arrestin-dependent pathways thereby not only delivering symptomatic relief in AD by improving memory and reducing anxiety, but also delivering neuroprotection that will maintain normal behaviour and extend life span.

## Results

### The M1-PD receptor is coupled normally to Gq/11 signalling, but is deficient in arrestin recruitment and internalisation

Our previous studies established that removal of all mass spectrometry (MS)-identified phosphorylation sites of the M1-receptor (22) together with all other putative serine/threonine phosphorylation sites in the third intracellular loop and C-terminal tail, generated a mutant receptor (M1-PD) that is normally coupled to Gq/11 signalling, but deficient in arrestin recruitment (17). Here, we have extended our previous studies by employing BRET biosensor assays to measure β-arrestin recruitment and receptor internalisation **(Figure 1A-D)**. In these experiments, HEK293 cells expressing the M1-PD receptor showed a reduction in acetylcholine-mediated β-arrestin recruitment and receptor internalisation compared to the wild-type receptor **(Figure 1C-D, Supplementary Figure 1A-D** and **Table 1)**. In contrast, coupling to the Gq/11 protein inositol phosphate pathway was near equivalent between M1-PD and wild-type receptors **(Figure 1E, Table 1)**.

**Figure 1.**
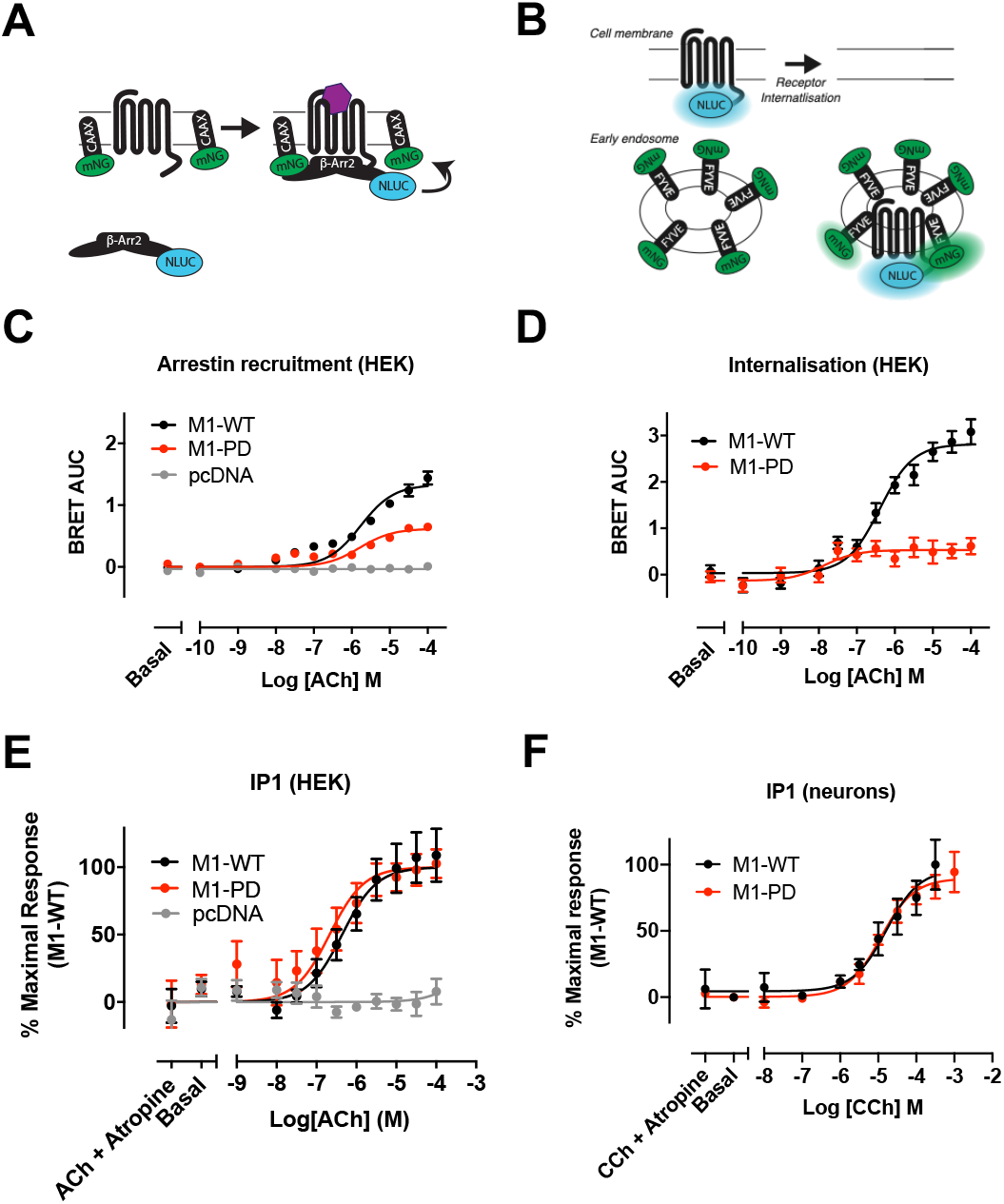
Arrestin recruitment and receptor internalisation of the M1-receptor is dependent on receptor phosphorylation. **(A-B)** Schematic of the bystander BRET assays for arrestin recruitment to the M1-receptor **(A)** and receptor translocation to early endosomes **(B). (C)** ACh-stimulated translocation of β-arrestin-2 to the cell membrane in HEK293T cells transfected with pcDNA3, M1-WT or M1-PD assessed by bystander BRET. Data are expressed as mean ± SEM of 5 independent experimented performed in triplicates. **(D)** Translocation of M1-WT and M1-PD to early endosomes in response to ACh treatment, assessed through a bystander BRET assay. Data are expressed as mean ± SEM of 5 independent experimented performed in triplicates. **(E)** IP1 accumulation after 60 minutes stimulation with ACh in HEK cells transiently transfected with the M1-WT and M1-PD constructs or the empty vector (pcDNA). Data are expressed as means ± S.E.M. of 4-7 independent experiments performed in duplicate or quadruplicate. **(F)** IP1 accumulation after 60 minutes stimulation with CCh in primary hippocampal-cortical neurons prepared from M1-WT or M1-PD mice. Data are expressed as means ± S.E.M. of 3 independent experiments performed in duplicate.

**Table 1.** M1-PD shows equivalent G protein-mediated responses compared to M1-WT, but impaired arrestin recruitment and receptor internalisation. Potency and maximum effect of agonist stimulated IP1 accumulation, β-arrestin-2 recruitment to the cell membrane or receptor translocation to early endosomes at M1-WT or M1-PD receptors. Agonists used for HEK cells and neurons were ACh or CCh, respectively. Data are expressed as the means ± S.E.M. of 3-7 independent experiments.

|  |  | **pEC<sub>50</sub>** | **E<sub>max</sub>** | **n** |
| --- | --- | --- | --- | --- |
| **IP1 (HEK)** | M1-WT | 6.4 ± 0.1 | 107.5 ± 7.3 | 7 |
|  | M1-PD | 6.7 ± 0.2 | 98.6 ± 7.3 | 5 |
| **Arrestin recruitment (HEK)** | M1-WT | 5.8 ± 0.1 | 100.0 ± 0.0 | 5 |
|  | M1-PD | 5.8 ± 0.1 | 46.6 ± 0.0 | 5 |
| **Internalisation (HEK)** | M1-WT | 6.4 ± 0.1 | 100.0 ± 0.0 | 5 |
|  | M1-PD | 7.9 ± 0.4 | 22.5 ± 0.0 | 5 |
| **IP1 (neurons)** | M1-WT | 4.8 ± 0.2 | 96.3 ± 9.5 | 3 |
|  | M1-PD | 4.9 ± 0.1 | 89.6 ± 4.3 | 3 |

We have previously described the generation of genetically engineered mice where the coding sequence for the M1-PD variant was knocked-in into the natural gene locus of the M1-receptor (chrm1) (17). In addition, to aid identification of receptor expression an haemagglutinin (HA) epitope tag was fused to the C-terminus of the M1-PD receptor. Control mice were also generated where the coding sequence for the mouse wild-type M1-receptor fused at the C-terminus with an HA-tag was similarly knocked into the M1-receptor locus (these mice are termed M1-WT mice). Consistent with previous studies (17), we show here that M1-PD transcription in the cortex and hippocampus of M1-PD mice is comparable to that of the M1-receptor in M1-WT mice **(Supplementary Figure 2A)**. Furthermore, quantification of receptor expression using Western blotting to detect the HA-tag revealed no significant (*p*>0.05) difference in receptor expression when comparing M1-WT and M1-PD mice (**Supplementary Figure 2B-C)**. Consistent with our findings in recombinant systems, we also show here that in primary hippocampal-cortical neurons prepared from M1-WT or M1-PD mice, agonist-stimulated inositol phosphate accumulation is equivalent (**Figure 1F, Table 1**).

### Prion-infected M1-PD mice show key hallmarks of disease earlier than M1-WT mice

Our previous work demonstrated that M1-receptor positive allosteric modulators (PAMs) can offer both symptomatic and disease modifying properties in a mouse model of terminal neurodegenerative disease (18). Mice inoculated with Rocky Mountain Laboratory (RML) prion-infected brain homogenate developed terminal neurodegenerative disease showing progressive neuronal loss, significant neuroinflammation and behavioural deficits. Here, we aimed to define the role of M1 receptor phosphorylation/arrestin-dependent signalling pathways in neurodegenerative disease progression. M1-WT or M1-PD mice were inoculated with control- (normal, healthy brain homogenate) or prion-infected brain homogenates, and receptor expression levels were found to be unchanged in control- and prion diseased mice at 16 weeks post inoculation (w.p.i.) **(Supplementary Figure 3A-C)**. Inoculation of mice with prion-infected brain homogenate induces accumulation of misfolded, insoluble prion protein that is resistant to digestion with proteinase K (PrPsc) (18, 21, 23). M1-PD mice inoculated with prion-infected brain homogenate show earlier increases in PrPsc accumulation in the hippocampus and cortex, compared to M1-WT mice with prion disease **(Figure 2A-B, Supplementary Figure 4A-B)**. Since the time-course of prion disease progression is dictated by the expression level of cellular prion protein (PrPc), whereby increased PrPc expression accelerates disease progression (21), we assessed PrPc expression in M1-PD mice. Importantly, we found that PrPc transcript levels were equivalent in M1-WT, M1-PD and M1-KO mice **(Supplementary Figure 5)**. These data show that prion disease had progressed faster in mice expressing a variant of the M1-receptor with reduced coupling to phosphorylation/arrestin signalling, suggesting that M1-receptor signalling through this pathway mediates a previously unappreciated neuroprotective effect.

**Figure 2.**
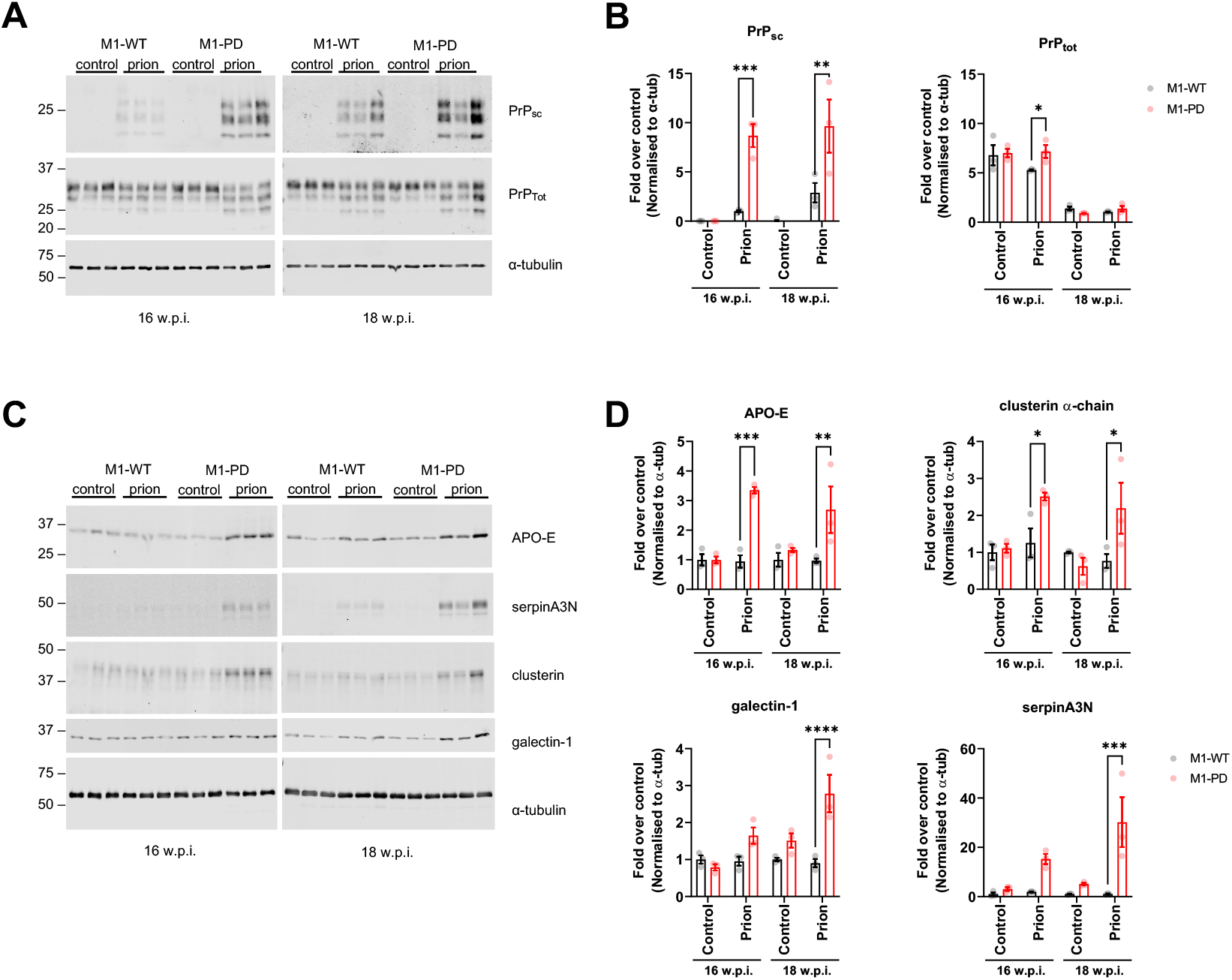
Prion infected M1-PD mice show accelerated appearance of disease markers in the hippocampus compared to M1-WT mice. Lysates were prepared from the hippocampus of control- or prion infected M1-WT and M1-PD mice at 16- and 18 w.p.i. and Western blot analysis was used to analyse the expression of a panel of pathological markers. **(A)** Lysates were incubated in the presence or absence of proteinase K prior to Western blot to detect non-digested scrapie prion protein (PrP_sc_) and total prion protein (PrP_tot_), respectively. Band analysis for PrP_sc_ and PrP_tot_ expression in **(B)** is shown as means ± S.E.M. of a ratio of α-tubulin expression.**(C)**Apolipoprotein-E (APO-E), serpinA3N, clusterin and galectin-1 were detected in the hippocampus and band analysis is shown in **(D)** as means ± S.E.M. of a ratio of α-tubulin expression relative to control-infected M1-WT (n=3 mice). All data were analysed using a two-way ANOVA with Sidak multiple comparisons where \**P*<0.05, \*\**P*<0.01, \*\*\**P*<0.001, \*\*\*\**P*<0.0001 (M1-WT vs. M1-PD).

We wanted to further test the possibility that uncoupling the M1-receptor from phosphorylation/arrestin signalling removed a neuroprotective component of receptor activity by monitoring biomarkers of disease severity. Recently, we have mapped the pathological changes in prion disease by conducting global transcriptomic and proteomic analyses of the hippocampus of prion diseased mice. We have found several protein markers previously associated with human AD that are significantly upregulated in prion disease. These included markers of neuroinflammation, glial fibrillary acidic protein (GFAP), clusterin, vimentin and galectin-1, and markers of adaptive responses to neurodegeneration including APO-E and the protease inhibitor serpinA3N (24). In this study we suggested that the upregulation of these proteins represent biomarkers of prion disease and indicators of disease severity (24). Here, we used the expression of these proteins to monitor disease progression in prion-infected M1-WT and M1-PD mice. Whereas M1-WT mice showed little change in the expression of APO-E, serpinA3N, clusterin α-chain and galectin-1 at 16- and 18 w.p.i., M1-PD mice showed a significant increase in all four of these prion disease biomarkers in the hippocampus and cortex **(Figure 2C-D and Supplementary Figure 4C-D)**. These results provide further evidence that the M1-PD variant of the M1-receptor allows accelerated prion-induced pathological changes.

### Prion-infected M1-PD mice show increased neuroinflammation compared to M1- WT mice

Murine prion disease, similarly to many human neurodegenerative diseases (25), is associated with profound neuroinflammation characterised by activation of both astrocytes and microglia (26). Here we further investigated the severity of prion disease in M1-WT and M1-PD mice by assessing the status of neuroinflammation. In prion diseased M1-PD mice, transcripts for GFAP and CD86, markers for astrocytes and microglia respectively, were significantly elevated compared to M1-WT mice **(Figure 3A, Supplementary Figure 6A)**. Furthermore, Western blotting for GFAP and vimentin, revealed a significant upregulation of astroglia in the hippocampus and cortex of prion-infected M1-PD mice **(Figure 3B-C, Supplementary Figure 6B-C)**. Importantly, the levels of both GFAP and vimentin were significantly higher in the hippocampus of RML-inoculated M1-PD mice in these time points compared to M1-WT mice **(Figure 3A-C)**. Protein level of GFAP and vimentin trended to higher levels in the cortex of M1-PD mice with prion compared to the respective wild-type animals at 18 w.p.i. **(Supplementary Figure 6B-C)**.

**Figure 3.**
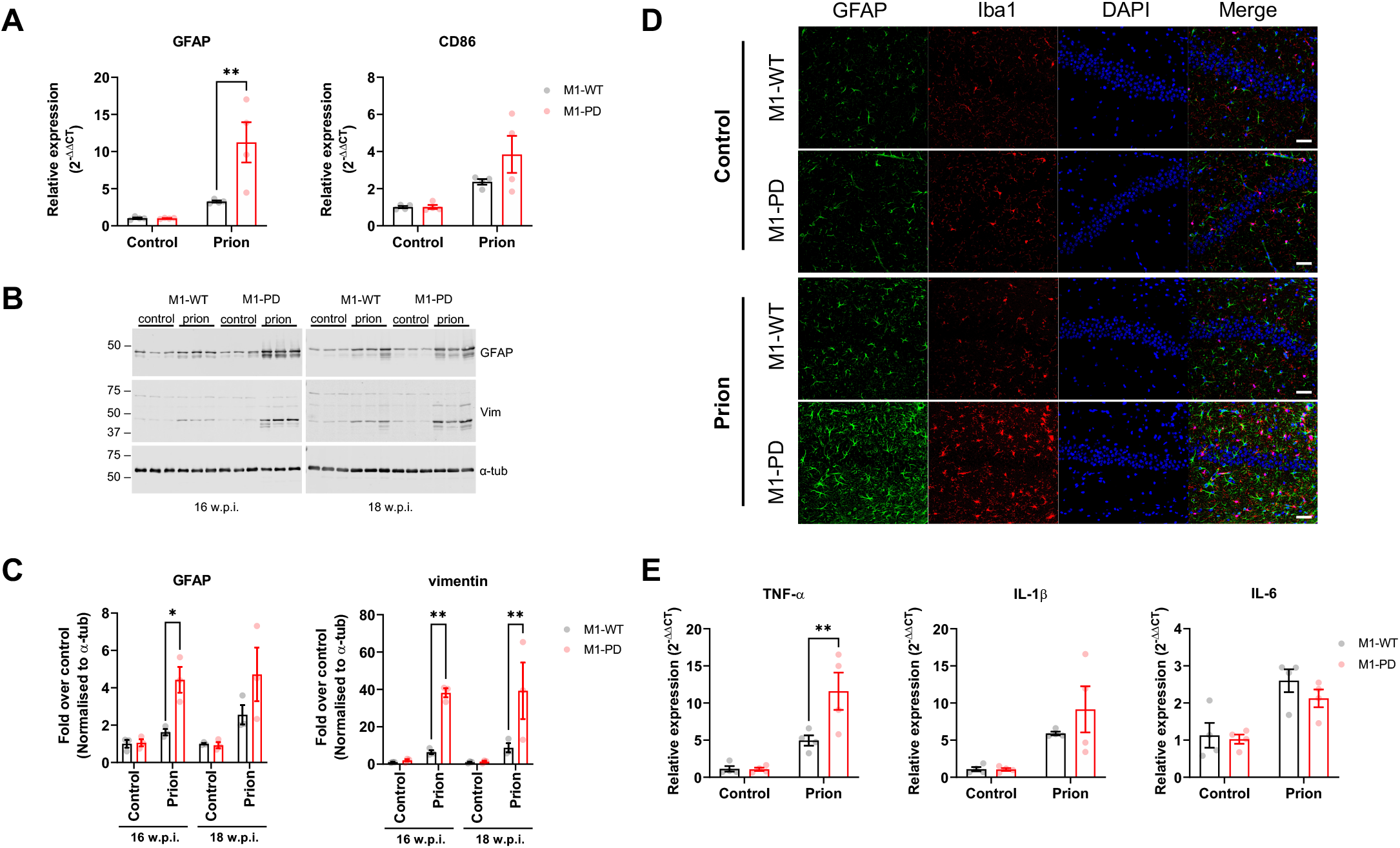
Neuroinflammation is exacerbated in the hippocampus of prion infected M1-PD mice compared to M1-WT controls. **(A)** mRNA levels of GFAP and CD86, markers of astrocytes and microglia respectively, were quantified using quantitative RT-PCR of hippocampus from control- or prion-diseased M1-WT or M1-PD mice at 16 weeks post inoculation (w.p.i.). Data is expressed as means ± S.E.M. of a ratio of α-tubulin RNA expression relative to M1-WT (n=4 mice). **P<0.01, two-way ANOVA with Sidak multiple comparisons (M1-WT vs. M1-PD). **(B-C)** Astrogliosis in the hippocampus was assessed using Western blot analysis of lysates prepared from control- or prion-infected mice at 16- and 18 w.p.i. Lysates were probed for astrocytic markers GFAP and vimentin (vim), and α-tubulin (α-tub) antibody was used as a loading control. **(C)** Band analysis for each blot was performed and data is shown as means ± S.E.M. of a ratio of α-tubulin relative to control M1-WT (n=3 mice). \**P*<0.05, \*\**P*<0.01, two-way ANOVA Sidak multiple comparisons (M1-WT vs. M1-PD). **(D)** Immunohistochemical staining for GFAP and Iba-1 in the hippocampus of control- and prion infected M1-WT and M1-PD mice at 16 w.p.i. The nuclei were stained blue with DAPI. Scale bar 100 μm. **(E)** Quantitative RT-PCR showing the expression of pro-inflammatory (TNF-α, IL-1β, IL-6) cytokines in the hippocampus of control- and prion infected M1-WT and M1-PD mice at 16 w.p.i. Data are expressed as a ratio of α-tubulin RNA expression relative to control M1-WT (n=4 mice). Data were analysed using two-way ANOVA with Sidak’s multiple comparisons, where \*\**P*<0.01 (M1-WT vs. M1-PD).

We further assessed the status of neuroinflammation by immunohistochemical staining of sections of the hippocampus and cortex. Staining for GFAP (astrocytes) and Iba-1 (microglia) demonstrated a profound increase in neuroinflammatory markers in the hippocampus and cortex of M1-PD, compared to M1-WT **(Figure 3D, Supplementary Figure 6D)**. Transcription of the pro-inflammatory cytokines TNF-α, IL-1β and IL-6 was elevated in the hippocampus and cortex of prion-infected M1-WT mice, compared to M1-WT mice inoculated with normal brain homogenate **(Figure 3E; Supplementary Figure 6E)**. Furthermore, we detected a significant increase in transcription of TNF-α in the hippocampus and cortex of M1-PD prion mice, compared to M1-WT, whereas IL-1β levels were elevated in the M1-PD versus M1-WT in the cortex only **(Figure 3E; Supplementary Figure 6E)**.

There were no differences in the expression of transcripts for anti-inflammatory cytokines, IL-4, IL-10, IL-11 and IL-13, in the cortex or hippocampus of prion-infected M1-WT or M1-PD mice **(Supplementary Figure 7)**. Importantly, expression of transcripts for GFAP, CD86 and the battery of pro- and anti-inflammatory cytokines tested above, were equivalent in non-infected M1-WT and M1-PD mice **(Supplementary Figure 8)**. Taken together, these data indicate that prion-infected M1-PD show increased neuroinflammation compared to M1-WT mice.

### M1-PD leads to earlier disease onset and shorter survival time

Prion-diseased mice exhibit behavioural deficiencies in burrowing activity, as reported previously (23), indicating a decline in hippocampal function. We assessed burrowing behaviour in prion-infected M1-WT and M1-PD mice and found that the M1-PD exhibited significant reductions in burrowing behaviour when compared to the M1-WT mice at 14- and 15 w.p.i **(Figure 4A)**. From 16 w.p.i. onwards, there were no significant differences in burrowing behaviour of prion-infected M1-WT and M1-PD mice. Importantly, the burrowing responses of M1-WT and M1-PD mice inoculated with control brain homogenate were equivalent at 9- and 17 w.p.i.

**Figure 4.**
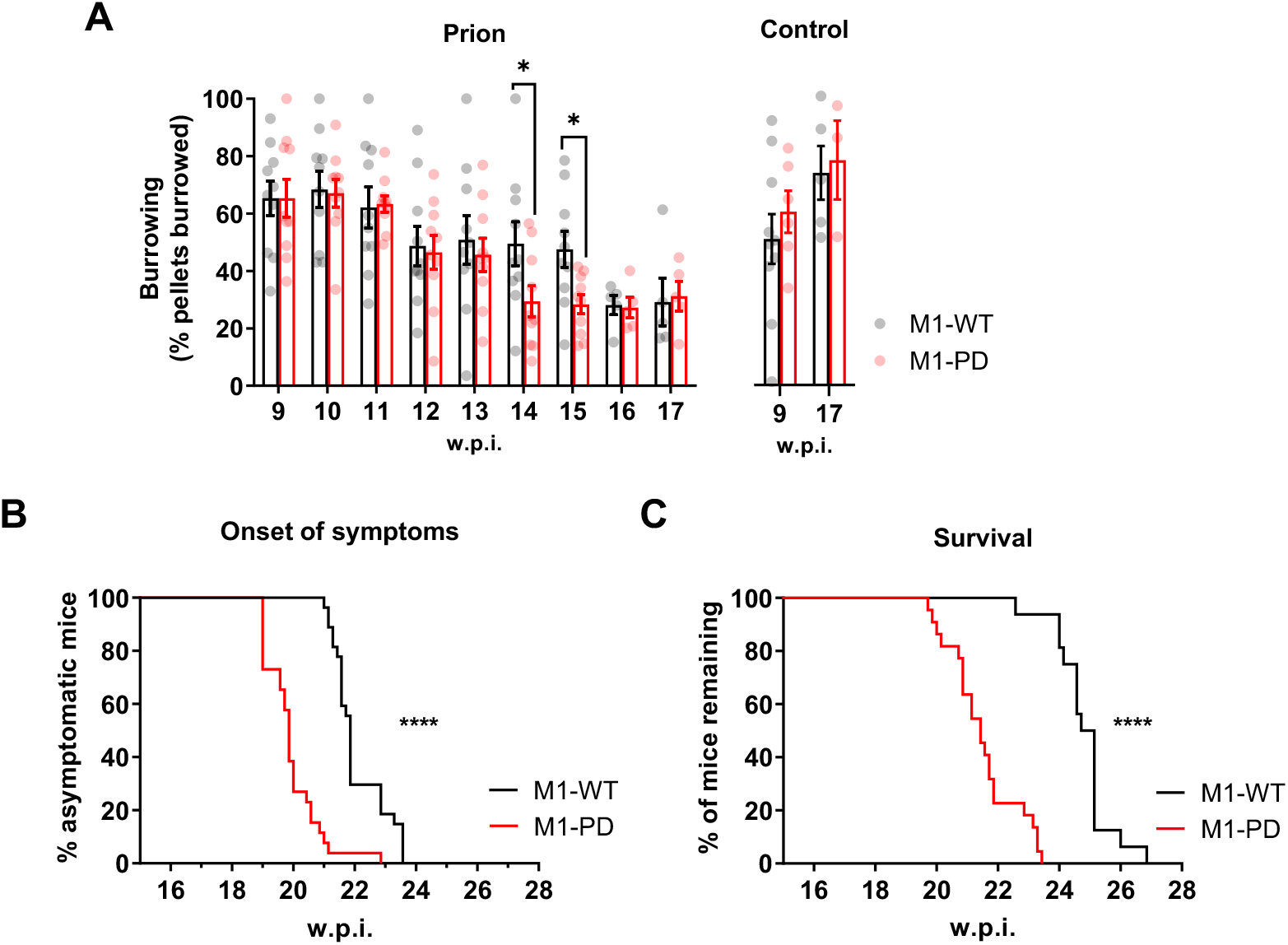
Removal of M1-receptor phosphorylation sites accelerates prion disease and decreases survival time. **(A)** Burrowing responses of control- or prion infected M1-WT and M1-PD mice were assessed from 9 weeks post inoculation (w.p.i.) (n=3-10 mice. \**P*<0.05; unpaired t test at each time point). Onset of at least two early indicators of prion disease (n=26-27) **(B)** and Kaplan-Meier survival plot (n=16-22) **(C)** for prion-infected M1-WT and M1-PD. Curves were analysed with a Gehan-Breslow-Wilcoxon test, where \*\*\*\**P*<0.0001.

Finally, we assessed the impact of M1-receptor phosphorylation on survival of prion diseased mice. Murine prion disease is a terminal neurodegenerative disease where clinical signs of disease – subdued behaviour, intermittent generalised tremor, erect penis, rigid tail, unsustained hunched posture, and mild loss of coordination – are evident from ∼21 w.p.i. in M1-WT mice **(Supplementary Figure 9)**. The appearance of at least two of the early indicator signs of prion disease occurred significantly earlier (P<0.0001) in M1-PD mice, compared to M1-WT **(Figure 4B)**. The median time for the onset of symptoms was 22 w.p.i. for M1-WT mice, and 20 w.p.i. for M1-PD mice. Mice expressing the phosphorylation-deficient M1-receptor mutant showed a significant (P<0.0001) acceleration of confirmatory scrapie diagnosis, with median time for terminal illness of 25 w.p.i. for M1-WT mice, and 21 w.p.i. for M1-PD mice **(Figure 4C)**. These data demonstrate that in addition to showing accelerated onset of behavioural abnormalities (burrowing), the M1-PD mice also showed an acceleration of the onset of clinical symptoms of prion disease.

## Discussion

By genetically engineering mice to express a phosphorylation-deficient variant of the M1-receptor we demonstrate here that physiological M1-receptor coupling to phosphorylation/arrestin-dependent pathways provides protection from prion-induced neurodegeneration. This was evident in the rapid onset of prion disease in mice expressing a variant of the M1-receptor (M1-PD) that has all potential phosphorylation sites removed (17). Previous studies from our laboratory have focused on the outcome of pharmacologically targeting the M1-receptor in prion disease. These earlier studies demonstrated that exogenous ligands that activate the M1-receptor can restore learning and memory deficits in prion disease and slow disease progression (18). Here we extend these studies by revealing an inherent, endogenously regulated, M1-receptor neuroprotective activity that results in suppression of markers of neurodegenerative disease and a reduction in neuroinflammation. Furthermore, our data show that this neuroprotective mechanism depends on receptor coupling to phosphorylation/arrestin-dependent mechanisms of signal transduction.

Our study has important implications for drug design. Like many members of the GPCR superfamily, muscarinic receptors mediate signal transduction in a bimodal fashion that involves both canonical G protein signalling and receptor phosphorylation/arrestin-dependent signalling. Whereas the molecular details of this phenomenon have been extensively studied in *in vitro* transfected cell systems, understanding the physiological importance of bimodal signalling and further, how this might have a pathophysiological impact, has been extremely challenging. We have approached this problem by genetically engineering mice to express variants of receptors where intracellular phosphorylation sites have been removed thereby reducing coupling to arrestin but maintaining coupling to heterotrimeric G proteins. This approach has been successfully employed, for example, in mapping physiological processes down-stream of the M3-receptor (27). Applying this same approach to the M1-receptor, a receptor widely considered as a validated target for improving memory loss and promising disease-modification in AD (7), our previous studies have established that activating ligands that maintain receptor phosphorylation/arrestin-dependent signalling can restore defective learning and memory in pre-clinical models of disease whilst minimising cholinergic adverse responses (17). Our study here extends these observations and suggests that M1-receptor ligands designed to promote receptor phosphorylation will have the additional benefit of driving neuroprotective receptor activity.

Our study also has important pathophysiological implications. AD is characterised by a loss of cholinergic neurons originating from the basal forebrain innervating the neocortex, amygdala, hippocampus and entorhinal cortex (28-30). The loss of cholinergic innervation is thought to be responsible for cognitive deficits in AD (29, 31), a hypothesis that underpins the use of cholinesterase inhibitors as the primary clinical strategy to improve memory loss in AD. The discovery here that M1-receptor activity has an inherent neuroprotective effect suggests that progressive loss of cholinergic neurons in AD will not only result in the loss of transmission in key memory centres but also a concurrent diminution of the M1-receptor neuroprotective effect. In this way, the loss of cholinergic neurons in AD with the associated reduction in acetylcholine-driven activation of post-synaptic M1-receptors might contribute to disease progression.

Global proteomic and transcriptomic studies have established that murine prion disease shows many of the key hallmarks of AD including neuroinflammation, mitochondrial dysfunction and oxidative stress (24). Furthermore, adaptive processes associated with the clearance of misfolded proteins commonly seen to be upregulated in AD are also seen to be upregulated in murine prion disease (24). These studies support the notion that neurodegenerative diseases propagated by the spread of *“prion-like”* misfolded proteins share common disease features (32). It is therefore significant that disease-markers common to AD and prion disease such as APOE and serpinA3N as well as markers of neuroinflammation are seen to be elevated in M1-PD prion infected mice. This indicates that the neuroprotection resulting from endogenous M1-receptor phosphorylation/arrestin-dependent signalling could be relevant to the protection against other neurodegenerative diseases that result from the accumulation of *“prion-like”* misfolded proteins.

In conclusion, we establish that the M1-receptor has inherent neuroprotective activity driven by receptor phosphorylation/arrestin-dependent signalling that results in the suppression of neuroinflammation and diminishes markers of neurodegenerative disease. Our data suggests that M1-receptor ligands that promote receptor phosphorylation/arrestin-dependent signalling will not only act to improve memory deficits in neurodegenerative diseases such as AD but will deliver neuroprotection that will maintain normal behaviour and extend life span.

## Materials and Methods

### Mouse maintenance and diet

Generation of the M1-WT, M1-PD and M1-KO strains used was described previously (17). All mice were bred as homozygous onto a C57Bl/6J background. Experiments in non-infected mice were conducted on male and female mice at 8-12 weeks old. Mice were fed ad libitum with a standard mouse chow. Animals were cared for in accordance with national guidelines on animal experimentation and all experiments were conducted under a UK Home Office project licence under the Animals (Scientific Procedures) Act (1986).

### RNA extraction

RNA extraction was performed immediately following tissue lysis using the Qiagen lipid tissue RNeasy Plus Mini kit as per manufacturer instructions. Briefly, following homogenisation in RLT Buffer containing 10% β-mercaptoethanol, the homogenate was centrifuged at 10,000 x g for 30 sec in a gDNA eliminator column. The aqueous phase containing RNA was collected and mixed with 70% ethanol, then applied to an RNeasy Mini spin column and subjected to centrifugation at 8000 x g for 15 sec at room temperature. The flow-through was discarded and the column was washed with guanidine containing stringent wash buffer, then washed with mild wash buffer, centrifuging for 2 min after the final wash to remove residual ethanol. RNA was eluted in 40 μl nuclease-free water.

### Determination of RNA concentration

RNA concentration was determined by measuring absorbance at 260 nm using a NanoDrop spectrophotometer. RNA purity was assessed using the A230/A260 and A260/A280 ratios, with ratios of 2–2.2 considered pure. RNA was stored at –80°C until use.

### qRT-PCR

For cDNA synthesis, 1 μg total RNA template per reaction was used using High Capacity cDNA Reverse Transcription Kit (Agilent). RNAase-free water (total 4.2 μl), 2 μl 10x RT buffer, 1 μl RT enzyme, 0.8 µl 25x dNTP Mix (100 mM) and 2µl 10x RT random primers were mixed together and incubated for 10 min at 25°C, followed by 120 min at 37°C, followed by 5 min at 85°C. Samples were then chilled at 4°C. Each reaction included control reactions in the absence of RT enzyme (-RT control). cDNA samples were stored at -20°C until qRT-PCR was performed. Each reaction was conducted in a total volume of 14 μl containing: 7 μl SYBR Green Master Mix, 1.4 μl primers (10 μM stock), 4.2 μl water and 1 μl cDNA (or –RT sample). Each reaction was performed in triplicate.

QuantiTect Primer Assays (Qiagen) were used for all the genes analysed. Sequences of the QuantiTect Primer Assays are not provided but the approximate location of primers within a specific gene can be viewed on the Product Detail of the manufacturer’s website.

| Primer Template | Primer assay name | Source and P/N |
| --- | --- | --- |
| CD86 (mouse) | Mm_Cd86_1_SG | (Qiagen, P/N: QT01055250) |
| Chrm1 (mouse) | Mm_Chrm1_1_SG | (Qiagen, P/N: QT00282527) |
| Gfap (mouse) | Mm_Gfap_1_SG | (Qiagen, P/N: QT00101143) |
| IL-1β (mouse) | Mm_Il1b_2_SG | (Qiagen, P/N: QT01048355) |
| IL-4 (mouse) | Mm_Il4_1_SG | (Qiagen, P/N: QT00160678) |
| IL-6 (mouse) | Mm_Il6_1_SG | (Qiagen, P/N: QT00098875) |
| IL-10 (mouse) | Mm_Il10_1_SG | (Qiagen, P/N: QT00106169) |
| IL-11 (mouse) | Mm_Il11_1_SG | (Qiagen, P/N: QT00122122) |
| IL-13 (mouse) | Mm_Il13_1_SG | (Qiagen, P/N: QT00099554) |
| Prion protein (mouse) | Mm_Prnp_1_SG | (Qiagen, P/N: QT00101080) |
| α-tubulin (mouse) | Mm_Tuba1b_1_SG | (Qiagen, P/N: QT00198877) |

### Sample preparation for Western blotting

Animal tissues were snap frozen in liquid nitrogen at sacrifice and stored at ™80°C. For preparation of tissue homogenates, frozen tissue was homogenised in ice-cold RIPA buffer (50 mM Tris-HCl pH 8, 150 mM NaCl, 0.5% (w/v) Na-deoxycholate, 1% (v/v) IGEPAL CA-630, 0.1% (v/v) SDS) including protease and phosphatase inhibitor cocktails (cOmplete™, Mini Protease Inhibitor Cocktail # 11836153001, cOmplete™, Mini Protease Inhibitor Cocktail # 11836153001, Sigma Aldrich -- Merck Life Science) using a sonicator at amplitude 3-5 Hz. The homogenates were incubated for 30 minutes at 4°C with end over end rotation and centrifuged at 21,000 x g for 15 minutes. Supernatants were transferred to fresh microcentrifuge tubes and stored at -80°C until use.

For the preparation of membrane extracts, frozen hippocampi and cortices were homogenised by sonication at 3-5 Hz amplitude in 500 µl of T/E buffer (10 mM Tris, 1 mM EDTA, pH 8.0) containing phosphatase and protease inhibitors. Samples were then centrifuged at 10,000 x g for 10 min at 4 °C. The supernatants were mixed with additional 500 µl T/E buffer, and then centrifuged at 15,000 x g for 1 hour at 4 °C. The pellets were then solubilised in 400 µL of RIPA buffer including phosphatase and protease inhibitors and incubated for at least 2 hours at 4°C with end over end rotation. After centrifugation of samples at 14,000 x g for 10 min at 4°C, the supernatants (membrane extracts) were transferred to fresh microcentrifuge tubes and stored at - 80°C until use.

Protein concentrations were determined by using the Micro BCA protein assay reagent kit according to the manufacturer’s instructions.

### Western blotting

Following preparation as described above, samples were incubated with Laemmli loading buffer containing 5% β-mercaptoethanol. Membrane extracts were incubated for 30 min at 37°C, whereas brain homogenates were heated at 95°C for 5 min, and briefly spun down before loading the gel. Samples (10 µg) were run in either 7.5% (>60 kDa), 12% (<60 kDa), -SDS-PAGE minigels at 60-150 V in Tris-glycine SDS running buffer (Serven biotech ltc, cat 20-6400-50). Nitrocellulose membranes were equilibrated in transfer buffer, before assembly the blot sandwich. The transfer was performed for 2 hours at constant 60V. Then, membranes were blocked for 30-45 min in 5% (w/v) fat-free milk in TBS-T (0.1% tween-20 in TBS at pH 7.4) at room temperature. Membranes were then incubated with the respective primary antibody at 4°C overnight or room temperature for 2 hours, and incubated with the respective secondary antibodies (1:10,000 for LI-COR antibodies) for one hour at room temperature. Proteins were visualised using LICOR Odyssey SA scanner using the appropriate lasers.

### Cell culture

HEK293T cells were maintained in Delbecco’s Modifeid Eagle Medium (DMEM) containing 10% FBS and maintained at 37°C and 5% CO_2_. HEK293T cells were transfected with polyethylenimine (PEI), using a 1:6 weight to weight ratio of DNA to PEI. 24 h post transfection cells were detached and seeded at either 20,000 cells/well into poly-D-lysine coated white 96-well plates for BRET assays, or 40,000 cells/well into poly-D-lysine coated clear 96-well plates for IP1 assay. Cells were then returned to the incubator and cultured for a further 24 h for BRET based assays or 48 h for IP1 accumulation assays.

### Primary neuronal culture

Tissue culture plates were coated using 4 µg/ml poly-D-lysine and 6 µg/ml Laminin Mouse Protein in DEPC-treated H_2_O and incubated overnight at 37 °C. Plates were then washed three times using DEPC-treated H_2_O and dried for 2h at room temperature.

The hippocampal and cortical areas of the brain were isolated from E16 mouse embryos. The tissues were chopped into smaller pieces, washed three times in Hanks’ balanced salt solution (HBSS), transferred to a 15 ml tube containing 4ml of TrypLE Select 10X and incubated at 37 °C for 10 min. TrypLE Select 10X was then inactivated by adding 8 ml of neurobasal complete media (Neurobasal Plus medium supplemented with 20 ml/L B-27 Plus, 0.292 g/L L-glutamine, 100 U/mL penicillin, 0.1 mg/L streptomycin) to the tubes followed by centrifugation at 200 x g for 5 min. The pellet was resuspended in neurobasal complete media to a final density of 5×10^5^ cells/ml. Cells were then seeded onto pre-coated plates and maintained at 37 °C in a 5% CO_2_ humidified atmosphere.

### IP1 Accumulation assay

HEK293T cells transiently transfected with vectors containing the M1-WT or M1-PD, or with the empty vector (pcDNA) were plated on 96-well plates at cell density of 40,000 cells/well for IP1 accumulation assays. Cells were washed and incubated in 1X stimulation buffer (Hank’s Balanced Salt Solution (HBSS) w/o phenol red containing 20 mM HEPES, 1.2 mM CaCl_2_; 30 mM LiCl; pH7.4) for one hour at 37°C prior drug treatments. 10X concentrated test compounds were added (5 μl/well) to the 96-well plates and incubated at 37°C for one hour. Following the treatment, the stimulation buffer was removed and lysis buffer (IP-One assay kit, CisBio) was added (40 µl/well). Following 10 minute-incubation on a shaker at 600 rpm, cell suspensions (7µl/well) were added to 384-well white proxiplates (PerkinElmer). The IP1-d2 conjugate and the anti-IP1 cryptate Tb conjugate (IP-One Tb™ assay kit, CisBio) were diluted together 1:40 in lysis buffer and 3 μl of the antibody mix were added to each well. The plate was incubated at 37°C for 1-24h and FRET between d2-conjugated IP1 (emission at 665 nm) and Lumi4™-Tb cryptate conjugated anti-IP1 antibody (emission at 620 nm) was detected using a CLARIOstar plate reader (BMG Labtech). Results were calculated from the 665/620 nm ratio and normalised to the maximum response stimulated by ACh.

Primary neuronal cells were seeded at a density of 5×10^4^ cells per well onto pre-coated 96 well plates and maintained at 37 °C in a 5% CO_2_ humidified atmosphere. On DIV7 cells were washed and incubated in stimulation buffer, and the assay was conducted as detailed above. Inhibition of agonist stimulation was measured by pre-incubating cells with 10 µM atropine for 30 min at 37 °C prior to addition of carbachol for 1 hour.

### BRET Arrestin assay

To assess arrestin recruitment to M1-WT and M1-PD, a bystander BRET assay was employed (33). HEK293T cells were co-transfected with plasmids encoding: 1) human β-Arrestin-2 fused to Nanoluciferase (Nluc) at its N-terminus; 2) the mNeonGreen (mNG) fluorescent protein fused with the prenylation CAAX sequence of KRas at its C-terminus; and 3) Either an empty pcDNA3 plasmid, a plasmid encoding wild-type mouse M1, or a plasmid encoding mouse M1-PD. For these transfections a DNA weight ratio of 1:25:5 was used for the Nluc-β-Arrestin-2 : mNG-CAAX : pcDNA3/M1-WT/M1-PD plasmids. After transfection, cells were cultured 24h, transferred to white 96-well plates and cultured a further 24h. For the assay, transfected cells in white 96-well plates were first washed twice with HBSS supplemented with 20mM HEPES (HBSS-H) and incubated for 30 min at 37°C. Nluc substrate, Coelenterazine 400a, was then added to a final concentration of 5 μM and incubated for 10 min. Dual 535 and 475 nm luminescent emission measurements were then taken at one minute intervals using a PherStar FS plate reader (BMG labtech) for 5 min prior to and 30 min following the addition of the indicated test compounds. Net BRET responses were calculated as the 535/475 ratio after correcting for both the well baseline and test compound vehicle response. BRET data was then reported as the area under the Net BRET curve for the 30 min following test compound addition.

### BRET Internalisation assay

Internalization of M1-WT and M1-PD was assessed using a bystander BRET assay designed to measure the translocation of M1 receptor to early endosomes (33). HEK293T cells were co-transfected with: 1) a plasmid encoding wild-type or PD mouse M1 fused at its C terminal to Nluc; and 2) a plasmid encoding mNG fused at its C terminal to the FYVE domain of endofin. A DNA weight transfection ratio of 1:2 was used for Nluc-tagged M1 receptor to mNG-FYVE. 24 h after transfection cells were transferred to white 96-well plates and then plates were cultured a further 24 h prior to the assay. For the assay, plates were washed twice with HBSS-H, before incubating in HBSS-H for 30 min at 37 °C. Coelenterazine 400a was added to a final concentration of 5 μM, plates were then incubated a further 10 min. BRET measurements were taken using a PherStar FS plate reader, measuring luminescent emission at 535 and 475 nm at 2 min intervals for 6 min prior to addition of test compounds, and a further 1 h after addition of test compounds. Net BRET responses were calculated as the 535/475 ratio after correcting for both the well baseline and test compound vehicle response. BRET data was then reported as the area under the Net BRET curve for the 60 min following test compound addition.

### Immunohistochemistry

Following heat-induced epitope retrieval, sections were washed in TBS + 0.1% triton x-100, and blocked overnight at 4°C in TBS, 0.1% triton X-100, 10% goat serum and 1% BSA. Incubation with primary antibodies (anti-GFAP, Sigma-Aldrich, P/N: G3893; anti-Iba1, Thermo Fisher, P/N: PA5-27436) was conducted in blocking buffer overnight at 4°C or for two hours at room temperature. Following three washes, slides were incubated with Alexa Fluor fluorescent secondary antibodies for 2 hours at room temperature in blocking buffer. Following three washes, slices were mounted on glass slides using VECTASHIELD HardSet Antifade Mounting Medium with DAPI, let dry overnight at 4°C and sealed using nail varnish. All images were taken using LSM 880 confocal laser scanning microscope (Zeiss).

### Prion infection of mice

Transgenic knock-in mice expressing HA-tagged M1-WT and M1-PD receptors were inoculated by intracerebral injection (under 3% isoflurane anasthesia) into the right parietal lobe with 1% brain homogenate infected with Rocky Mountain Laboratory (RML) prion aged 3 to 4 weeks as described previously (18). Control mice received 1% normal brain homogenate.

### Burrowing

Assessment of burrowing on control and prion-infected M1-WT or M1-PD mice was conducted from 9 w.p.i. The burrowing test involved mice being placed into individual cages (22×36 cm) with a plastic cylinder filled with 140g of food pellets. Food remaining in the cylinders after 2 hours was weighed and the amount displaced (“burrowed”) was calculated. Prior to the burrowing test, mice were placed in the burrowing cage for a 2-hour period. The test was then repeated on a weekly basis.

### Symptoms scoring and survival analysis

Prion-infected mice were scored according to the appearance of recognised early indicator and confirmatory signs of prion disease. Early indicator signs included piloerection, sustained erect ears, erect penis, clasping of hind legs when lifted by tail, rigid tail, unsustained hunched posture, mild loss of coordination or being subdued. Confirmatory signs of prion disease included ataxia, impairment of a righting reflex, dragging of limbs, sustained hunched posture and/or significant abnormal breathing. Survival times were calculated based on the presence of 2 early indicator signs plus 1 confirmatory sign, or 2 confirmatory signs. At this time, mice would be humanely killed.

## Acknowledgements

This work is partially funded by an MRC Industrial CASE studentship (MR/P016693/1; MS), a University of Glasgow Lord Kelvin Adam Smith Fellowship (SJB), an MRC MICA (MR/P019366/1; ABT, SJB), a Wellcome Trust Collaborative Award (201529/Z/16/Z; ABT), an Academy of Medical Sciences Springboard Award (SBF004\1033; BDH) and an Alzheimer’s Research UK (ARUK) David Hague Early Career Investigator of the Year Award (ARUK-YI2020-002; SJB). We acknowledge the BSU facilities at the Cancer Research UK Beatson Institute (C596/A17196) and the Biological Services at the University of Glasgow.

## Notes

### Competing Interest Statement

The authors have declared no competing interest.

