## Supplementary figures for "Phosphorylation of the M1 muscarinic acetylcholine receptor mediates protection in neurodegenerative disease"

**Supplementary data**

**Supplementary Figure 1 Arrestin recruitment and receptor internalisation is impaired when phosphorylation sites are removed from the M1-receptor. (A-B)** Time-course of ACh-stimulated translocation of β-arrestin-2 to the cell membrane in HEK293T cells transfected with M1-WT **(A)** or M1-PD **(B)** assessed by bystander BRET. Data are expressed as mean ± SEM of 5 independent experimented performed in triplicates. **(C-D)** Time course of translocation of M1-WT **(C)** and M1-PD **(D)** to early endosomes in response to ACh treatment, assessed through a bystander BRET assay. Data are expressed as mean ± SEM of 5 independent experimented performed in triplicates.

**Supplementary Figure 2 M1 receptors are equivalently expressed in M1-WT and M1-PD mice. (A)** Quantitative analysis of M1-receptor RNA expression in M1-WT, M1-PD and M1-KO cortex and hippocampus. Data is expressed as means ± S.E.M of a ratio of α-tubulin RNA expression relative to M1-WT (n=4 mice). **(B)** Western blot analysis of solubilsed membranes from M1-WT, M1-PD and M1-KO cortex and hippocampus for detection of HA-tagged receptors. Data shown are from three separate mice for the M1-WT and M1-PD genotypes. $Na^+/K^+$ ATPase expression was used as a loading control. **(C)** Bands from (**B**) were analysed for (semi)quantification of receptor expression and data is expressed as a ratio of $Na^+/K^+$ ATPase expression relative to the M1-WT (n=3 mice).

**Supplementary Figure 3 Control- and prion infected M1-PD mice show equivalent receptor expression and signalling compared to M1-WT mice. (A)**

Quantitative analysis of M1-receptor RNA expression in the cortex and hippocampus of 16 weeks post-inoculation (w.p.i.) control- and prion-infected, M1-WT and M1-PD mice. Data is shown as means ± S.E.M of a ratio of α-tubulin RNA expression relative to M1-WT (n=4 mice). **(B)** Solubilised membranes from 16 w.p.i. control- and prion-infected M1-WT and M1-PD cortex and hippocampus were probed in a Western blot for the expression of M1-receptor using an antibody against the HA tag. Data shown are two separate mice for the control- and prion-infected M1-WT and M1-PD genotypes. M1-KO was also probed as negative control, and $Na^+/K^+$ ATPase expression was used as a loading control. **(C)** Band analysis from the western blot **(B)** was conducted for (semi)quantification of receptor expression and data is expressed as means ± S.E.M. of ratios of $Na^+/K^+$ ATPase expression relative to the M1-WT (n=2 mice).

**Supplementary Figure 4 Prion infected M1-PD mice show accelerated appearance of disease markers in the cortex compared to M1-WT mice.** Lysates were prepared from the cortex of control- or prion infected M1-WT and M1-PD mice at 16- and 18 w.p.i. and Western blot analysis was used to analyse the expression of a panel of pathological markers. **(A)** Lysates were incubated in the presence or absence of proteinase K prior to Western blot to detect non-digested scrapie prion protein ($PrP_{sc}$) and total prion protein ($PrP_{tot}$), respectively. Band analysis for $PrP_{sc}$ and $PrP_{tot}$ expression in **(B)** is shown as means ± S.E.M. of a ratio of α-tubulin expression (n=3). **(C)** Apolipoprotein-E (APO-E), serpinA3N, clusterin and galectin-1 were detected in the cortex and band analysis is shown in **(D)** as means ± S.E.M. of a ratio of α-tubulin expression relative to control-infected M1-WT (n=3 mice). Data were analysed using a two-way ANOVA with Sidak multiple comparisons where **$P$<0.01, ****$P$<0.0001 (M1-WT vs. M1-PD).

**Supplementary Figure 5 Prion protein (PrP) mRNA expression is unchanged in the hippocampus and cortex of M1-PD mice compared to M1-WT.** Quantitative RT-PCR showing the expression of PrP RNA in the cortex and hippocampus of M1-WT and M1-PD mice. Data are expressed as a ratio of α-tubulin RNA expression (n=4 mice).

**Supplementary Figure 6 Neuroinflammation is exacerbated in the cortex of prion infected M1-PD mice compared to M1-WT controls. (A)** mRNA levels of GFAP and CD86, markers of astrocytes and microglia respectively, were quantified using quantitative RT-PCR of cortex from control- or prion-diseased M1-WT or M1-PD mice at 16 weeks post inoculation (w.p.i.). Data is expressed as means ± S.E.M. of a ratio of α-tubulin RNA expression relative to M1-WT (n=4 mice). **(B-C)** Astrogliosis in the cortex was assessed using Western blot analysis of lysates prepared from control- or prion-infected mice at 16- and 18 w.p.i. Lysates were probed for astrocytic markers GFAP and vimentin (vim), and α-tubulin (α-tub) antibody was used as a loading control. **(C)** Band analysis for each blot was performed and data is shown as means ± S.E.M. of a ratio of α-tubulin relative to control M1-WT (n=3 mice). *$P$<0.05, two-way ANOVA Sidak multiple comparisons (M1-WT vs. M1-PD). **(D)** Immunohistochemical staining for GFAP and Iba-1 in the cortex of control- and prion infected M1-WT and M1-PD mice at 16 w.p.i. The nuclei were stained blue with DAPI. Scale bar 100 μm. **(E)** Quantitative RT-PCR showing the expression of pro-inflammatory (TNF-α, IL-1β, IL-6) cytokines in the cortex of control- and prion infected M1-WT and M1-PD mice at 16 w.p.i. Data are expressed as a ratio of α-tubulin RNA expression relative to control M1-WT (n=4 mice). Data were analysed using two-way ANOVA with Sidak multiple comparisons, where $^*P<0.05$, $^{**}P<0.01$ (M1-WT vs. M1-PD).

**Supplementary Figure 7 Expression of anti-inflammatory cytokines in control- and prion infected M1-WT and M1-PD mice.** Quantitative RT-PCR showing the expression of anti-inflammatory cytokines, **(A-B)** IL-4, **(C-D)** IL10, **(E-F)** IL-11 and **(G-H)** IL-13 in the cortex **(A, C, E, G)** and hippocampus **(B, D, F, H)** of control- or prion diseased M1-WT or M1-PD mice at 16 w.p.i. Data are expressed as a ratio of α-tubulin RNA expression (n=4 mice).

**Supplementary Figure 8 Expression of neuroinflammatory markers and cytokines in M1-WT and M1-PD mice. (A-B)** Quantitative RT-PCR showing the expression of **(A)** GFAP and **(B)** CD86, markers of astrocytes and microglia respectively, in the cortex or hippocampus of M1-WT or M1-PD mice. **(C-I)** Quantitative RT-PCR showing the expression of pro-inflammatory cytokines **(C)** TNF-α, **(D)** IL-1β and **(E)** IL-6, and anti-inflammatory cytokines **(F)** IL-4, **(G)** IL-10, **(H)** IL-11 and **(I)** IL-13 in the cortex or hippocampus of M1-WT or M1-PD mice. Data are expressed as a ratio of α-tubulin RNA expression (n=4 mice).

**Supplementary Figure 9 Removal of M1-receptor phosphorylation sites accelerates the onset of prion disease indicators.** Symptomatic mice were analysed according to the appearance of recognised early disease indicators including **(A)** subdued (n=23), **(B)** intermittent generalised tremor (n=8-15), **(C)** erect penis (n=5-6), **(D)** rigid tail (n=16-19), **(E)** unsustained hunched posture (n=13-14), and **(F)** mild loss of coordination (n=11-12). Curves were analysed with a Gehan-Breslow-Wilcoxon test, where $^{**}P < 0.01$; $^{****}P < 0.0001$.

Supplementary Figure 1

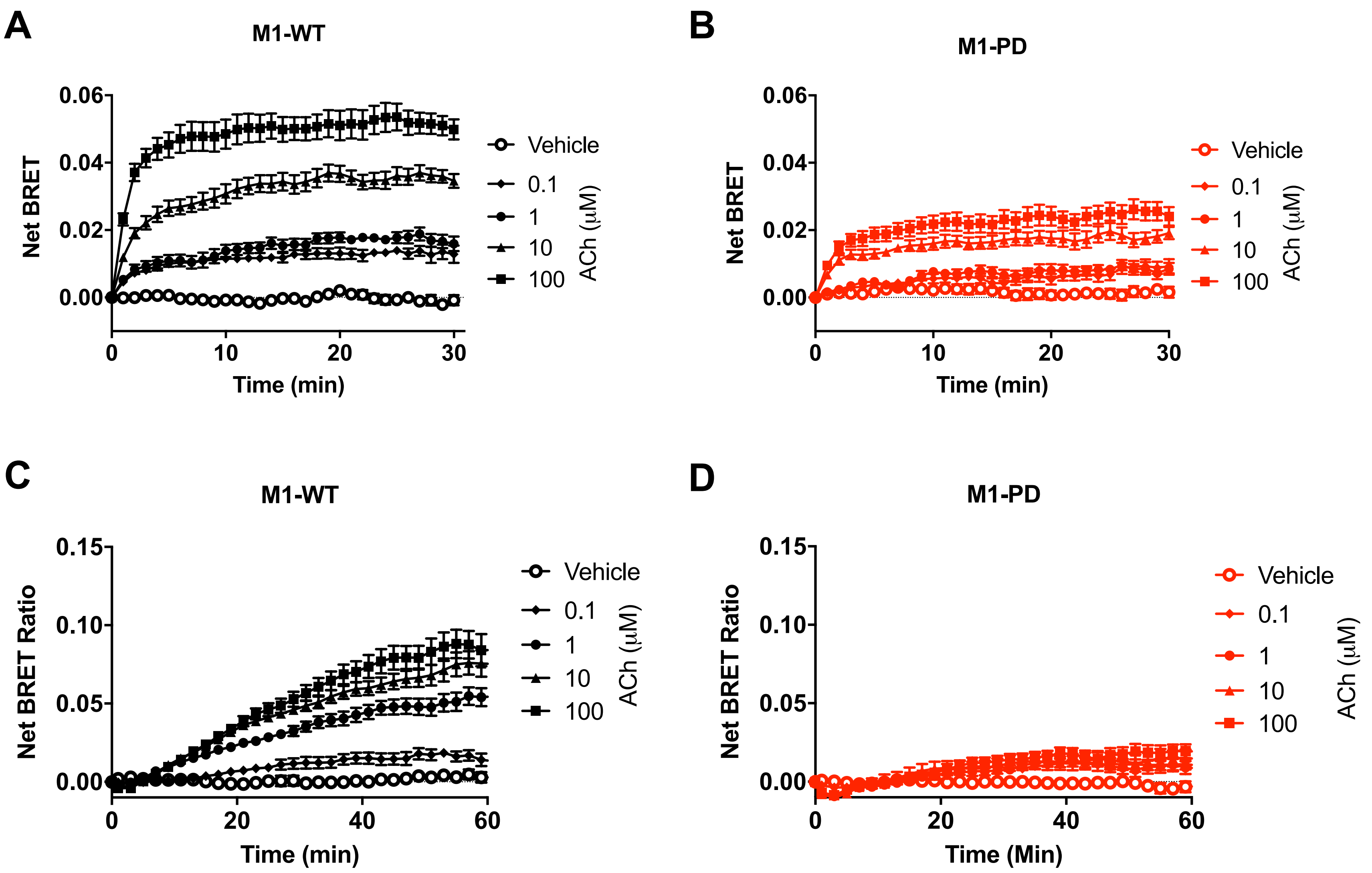

### Supplementary Figure 2

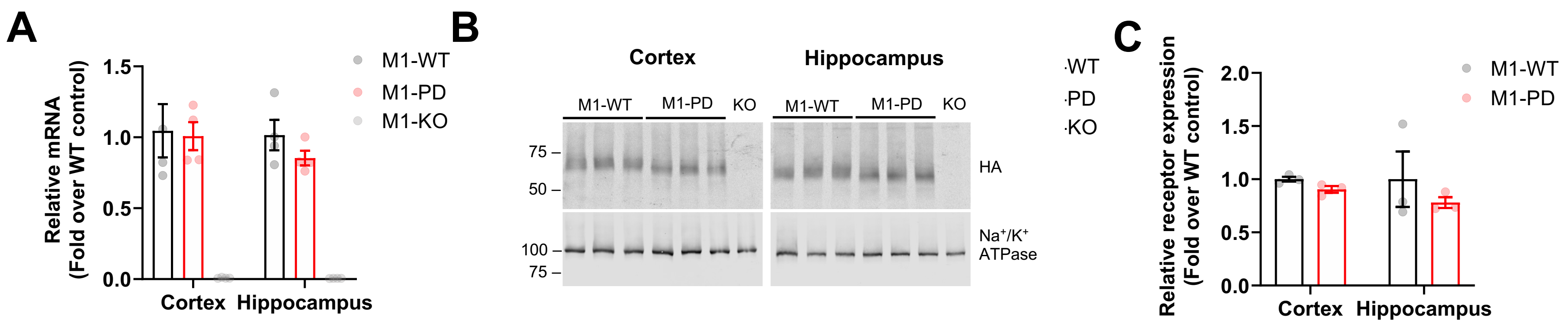

### Supplementary Figure 3

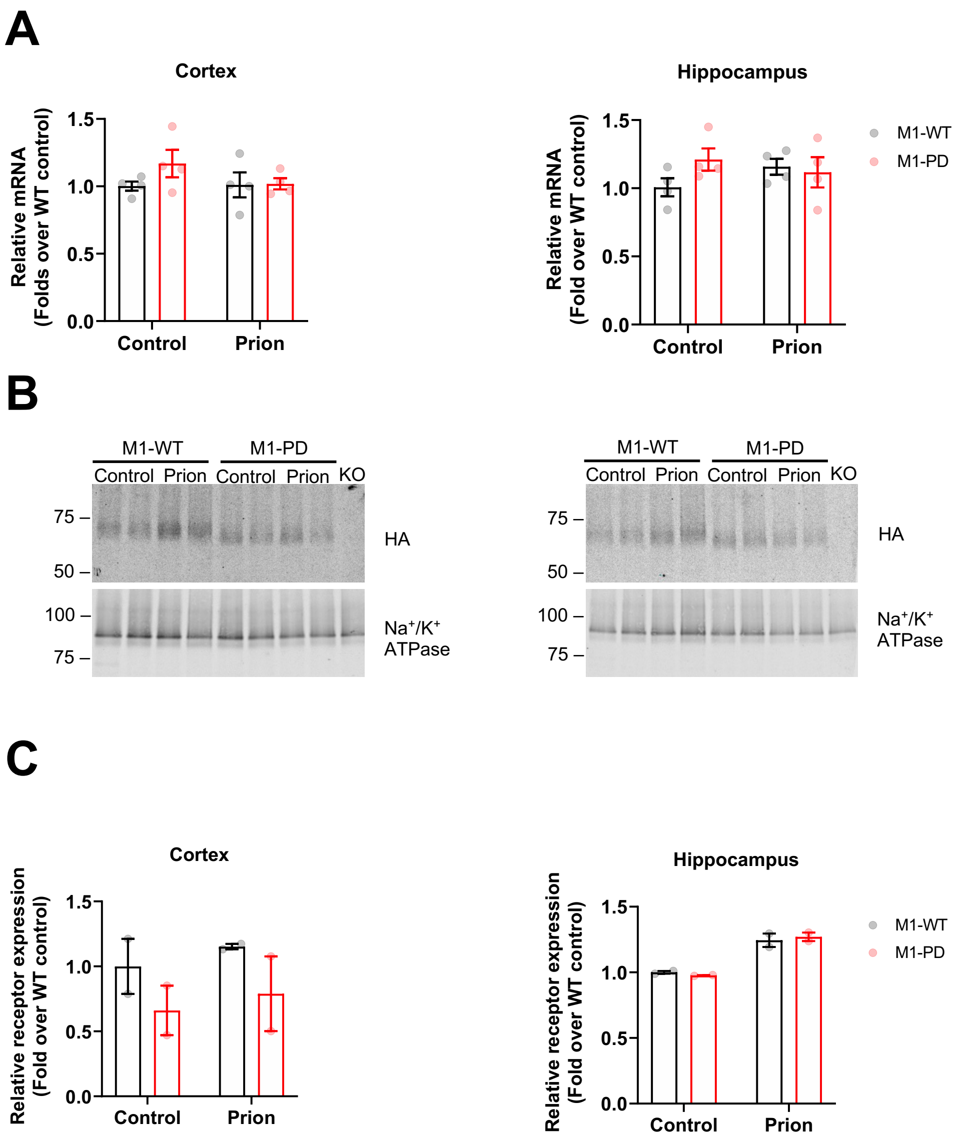

### Supplementary Figure 4

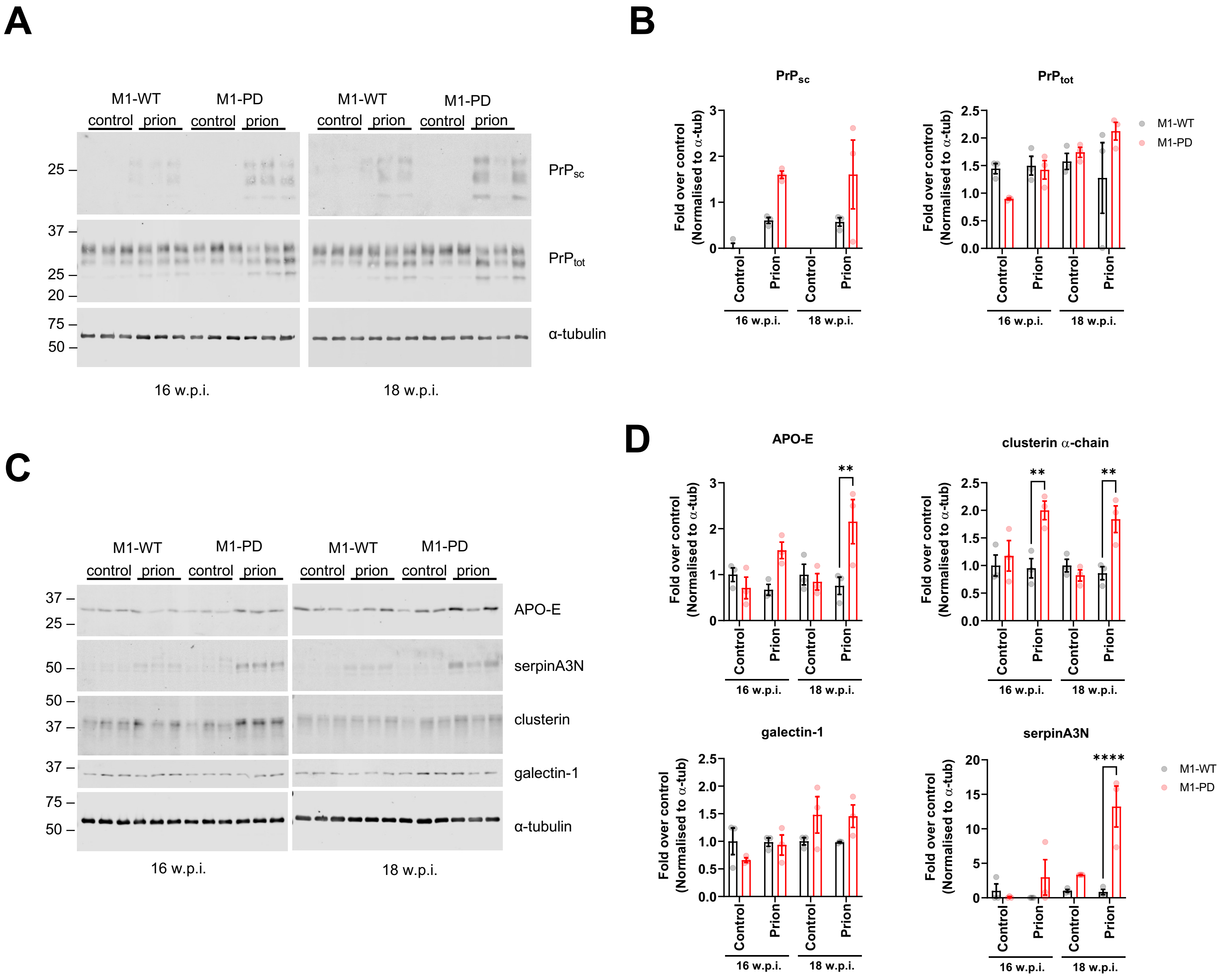

Supplementary Figure 5

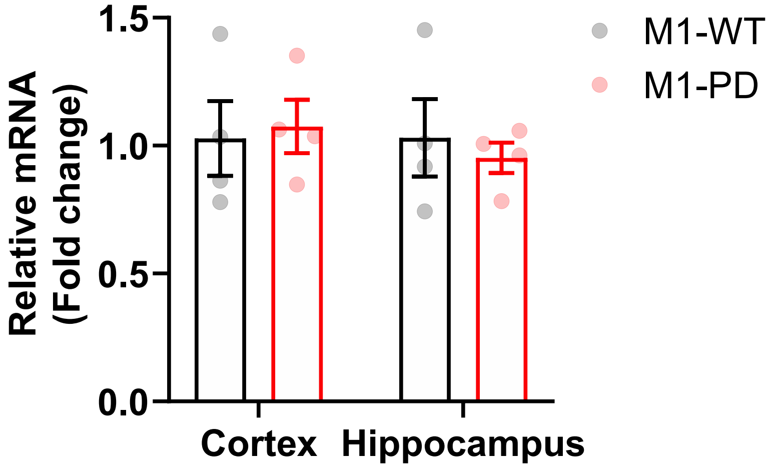

### Supplementary Figure 6

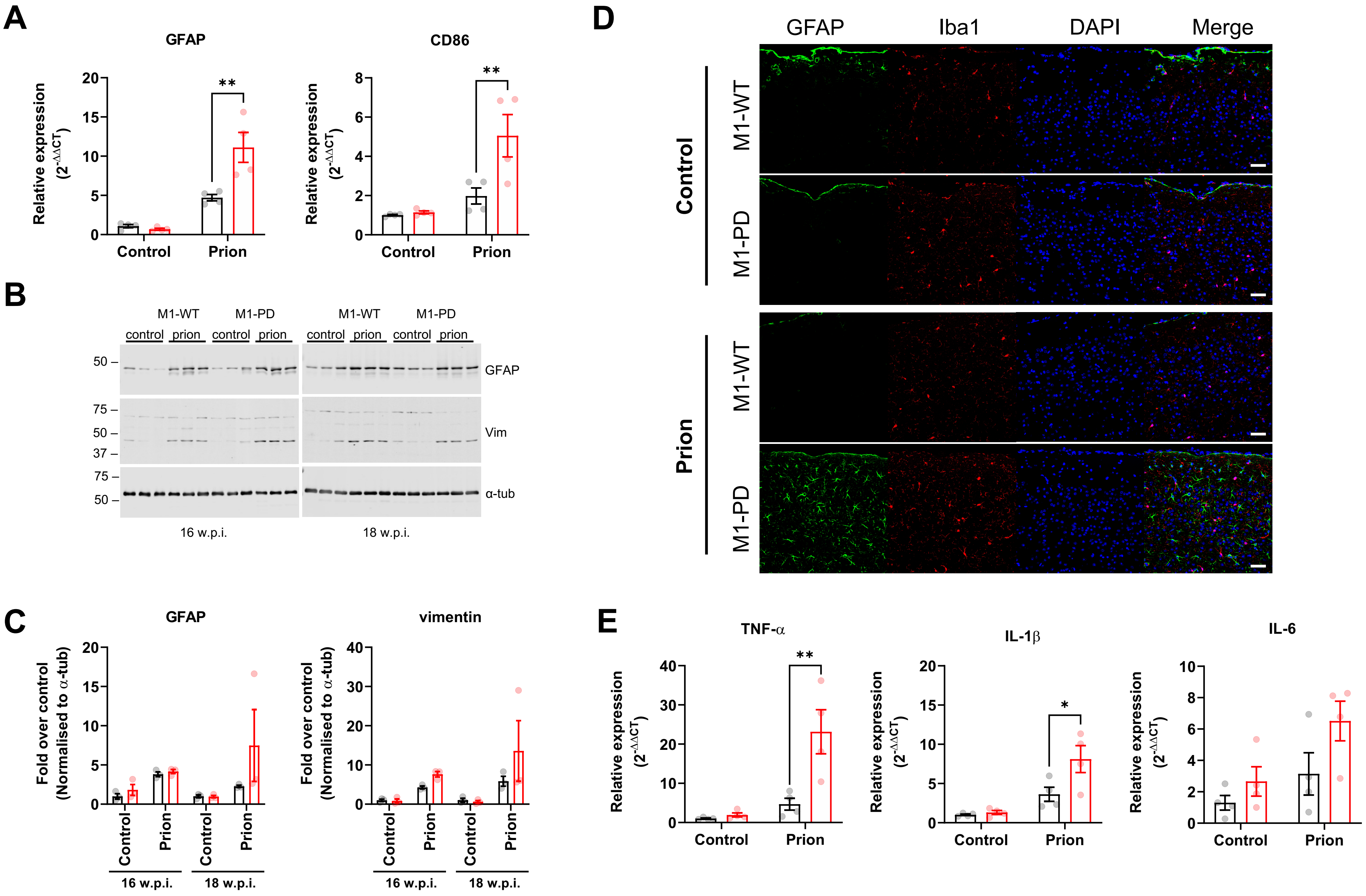

Supplementary Figure 7

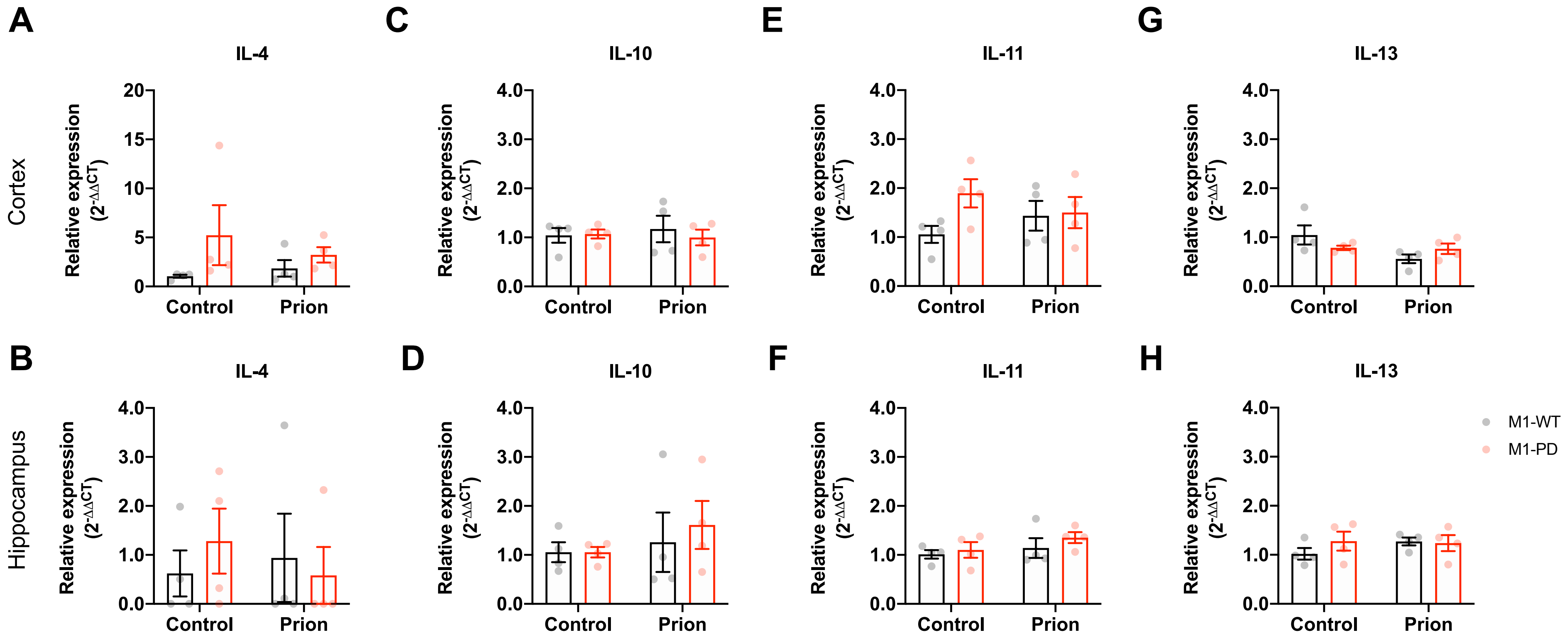

### Supplementary Figure 8

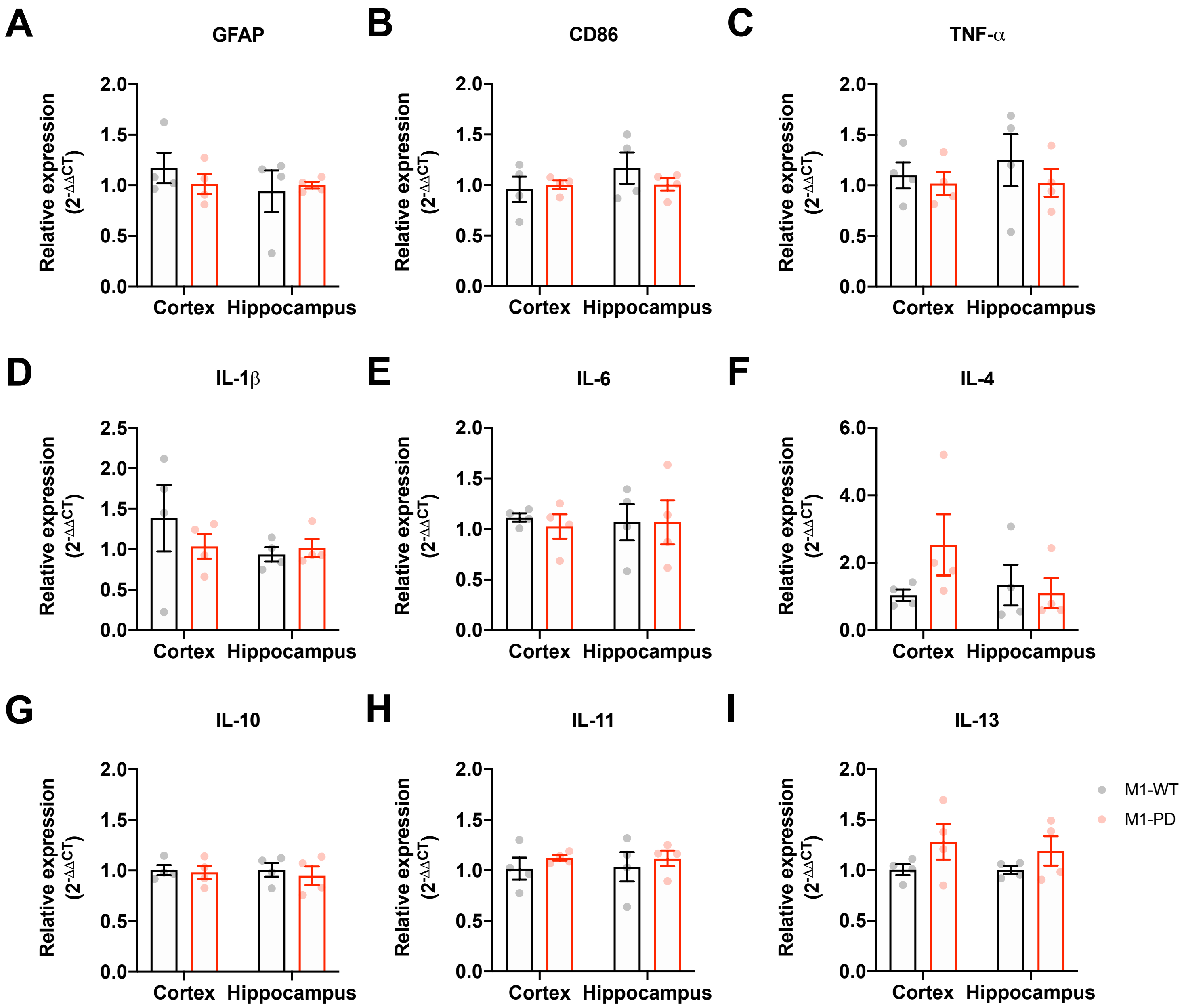

Supplementary Figure 9

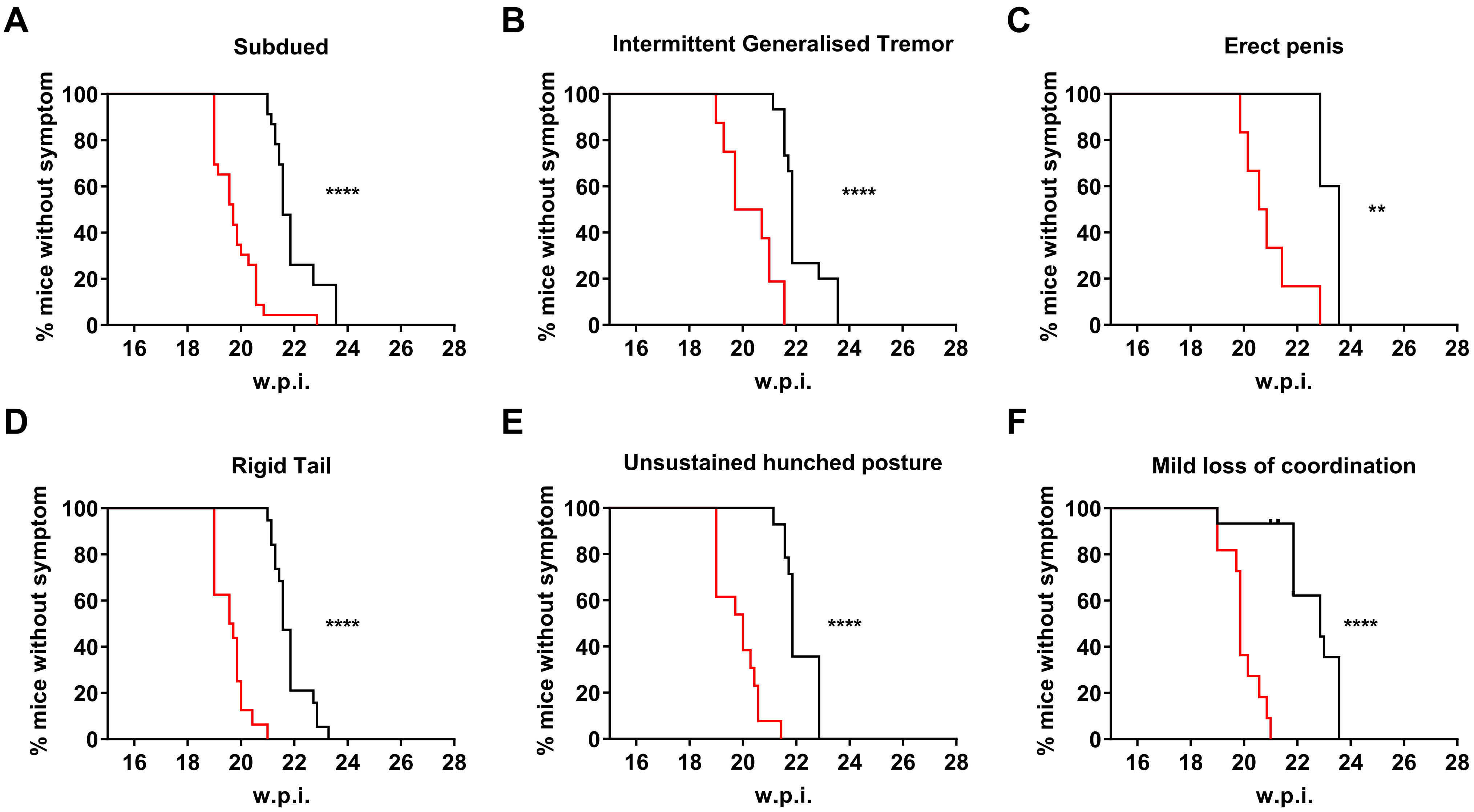
